# A novel PS4 criterion approach based on symptoms of rare diseases and in-house frequency data in a Bayesian framework

**DOI:** 10.1101/2020.07.22.215426

**Authors:** You Kyung Cho, Dhong-gun Won, Changwon Keum, Beom Hee Lee, Go Hun Seo, Byung-Chul Lee

## Abstract

The American College of Medical Genetics (ACMG) and Genomics/Association for Molecular Pathology (AMP) previously reported standardized guidance for the assessment of genetic variants. One of the criteria regarding the prevalence in a case-control study, PS4, is important due to its evidence of pathogenicity. Despite recent studies approaching gene- and disease-specific probands, interpretation of a variant to PS4 still has certain limitations for rare variants. Here, we suggest a generalized method, Bayesian odds ratio (BayesianOR), applicable to PS4 via decomposing a disease to its symptoms and applying a Bayesian framework. Using this approach, we demonstrate reproducibility of the calculation of the original odds ratio from well-studied epilepsy data and verify the applicability to in-house frequencies for various rare diseases. In addition, BayesianOR showed a significant difference in tendency with different ClinVar pathogenicity, using in-house data. Thus, the novel method described here should provide an improved interpretation of sequence variants. Furthermore, we anticipate that it will enhance the diagnosis of patients with rare diseases.

## Introduction

With advances in molecular testing technology, such as whole exome sequencing (WES) and whole genome sequencing (WGS), the number of genes associated with a genetic disorder and novel gene sequence variants is increasing rapidly. The American College of Medical Genetics and Genomics (ACMG) and the Association for Molecular Pathology (AMP) published standards and guidelines for the interpretation of sequence variants in 2015^1^. These guidelines classify variants as “pathogenic”, “likely pathogenic”, “uncertain significance”, “likely benign”, or “benign”, based on a combination of criteria with different levels of evidence defined as very strong, strong, moderate, and supporting, all contributing to an accurate genetic diagnosis and one that is consistent among laboratories. Each criterion is categorized by population frequency data, variant type and location, case-level data, functional and computational data, and reputable sources^2^.

In particular, case-level data include information regarding patient phenotypes related to variants, other detected variants of interest, patient family data, and whether the variant has been detected in unrelated affected individuals. PS4 is a criterion that the prevalence of the variant in affected individuals is significantly increased compared with that of controls, in a case-control study with an odds ratio (OR) > 5.0 and confidence interval not including 1.0^1^. However, case-control study data are not easily available for rare genetic diseases. The ACMG/AMP guidelines have suggested another way to evaluate variants such that the same variant observed in multiple unrelated affected individuals with consistent phenotypes, and its absence in population databases, could be used as PS4 evidence at a moderate level of evidence^1^.

Recently, the Clinical Genome Resource (ClinGen) published reports regarding variant interpretation according to ACMG guidelines, focusing on unique features of particular genes or genomic regions^3–8^. These reports describe the number of observations in unrelated affected individuals and any of modified strengths of each rule, according to disease-gene specifications^3–8^. These observations can be acquired through internal lab data, publicly available data, and through scientific literature. However, representative public data, such as ClinVar, does not provide detailed information regarding the count of unrelated affected individuals and their phenotype^9^. In addition, there might be a possibility that one proband reported multiple times in the literature, public database, or in a clinical testing cohort, may result in duplicate counting. It is a time-consuming and labor-intensive process to count unrelated affected individuals because of the difficulty in extracting this information automatically. Moreover, there remain certain hurdles to the general application of such criteria to all rare genetic disorders. Herein, we report a novel approach of expanded PS4 criterion applicable to a broad range of rare diseases.

## Materials and Methods

The PS4 criterion for rare variants is not applicable in many cases due to a lack of positive cases of the corresponding disease. To overcome this limitation, we propose a method of using symptoms instead of the disease itself. The basic assumption of this approach is that a disease can be decomposed into multiple symptoms. Since observed cases of each symptom can be accumulated by sharing symptoms among diseases (Supplement figure 1), the calculation of OR for some rare variants becomes possible, and will provide greater discriminative prevalence between the affected and general population for such rare variants.

### Symptom-based OR calculation using a Bayesian framework

According to the original ACMG/AMP guidelines, to determine whether a discovered variant is classified as a PS4 criterion or not, the OR should be obtained in a case-control study^1^. The odds ratio formula is,

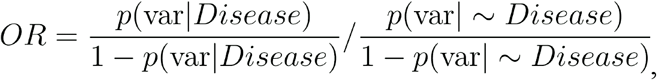

where var is the variant of interest.

Expanding the above equation to the symptom-based concept, the Bayesian odds ratio (BayesianOR) is defined as the disease replaced with symptom set *S*, which constitutes the disease, and gnomAD for the Control Group^10^. The formula can be represented as,

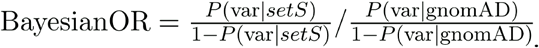

*P*(var|*setS*) can changed to *P*(*setS*|var)*P*(var)/*P*(*setS*) using the Bayesian rule. We assume that the probability of set *S* can be inferred by an additive model of summation of each probability of a symptom given the variant. Then, *P*(*setS*|var) can be

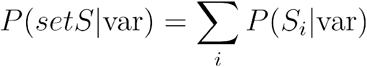

where *S_i_* is an element of set *S*.

To utilize internal data, *P*(*S_i_*|var) should be derived from *P*(var|*S_i_*) because *P*(*S_i_*|var) cannot be obtained from in-house data due to the number of variants from the affected patients. Further, by applying the Bayesian rule, *P*(*S_i_*|var) can be substituted to

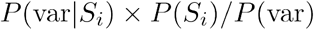

Thus, the final formula for the calculation of BayesianOR is,

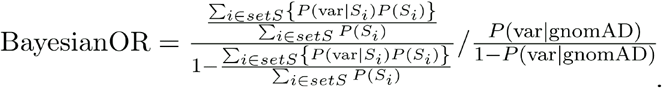

To calculate the BayesianOR score, prior prevalence of symptoms should also be calculated. We define the equation as follows, 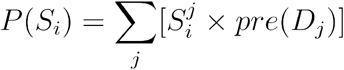 where 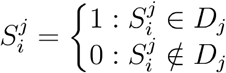 and *pre*(*D_j_*) is the prevalence of a disease *j*.

Since the basic assumption is that each disease is a union of symptoms belonging to the disease, an anonymous rare disease *D_i_* can be depicted as,

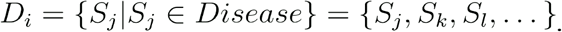

The probabilities of intersection among rare diseases are ignored based on the assumption that *P*(*S_i_* ∩ *S_j_*) are extremely rare. In the case that either P(var) or P(var|S_i_) is zero, the minimum probabilities are inferred as 1.0e^-1^ and 2.5e^-7^, respectively,^11^.

### Data preparation

#### Sample collection

This research was approved by the Institutional Review Board for Human Research of the Asan Medical Center (IRB numbers: 2018-0574 and 2018-0180). All samples were collected from 6 major domestic hospitals from March 2018 to December 2019, and the symptoms of each patient were described by physicians through the 3billion website (http://portal.3billion.io/). Blood, saliva, or buccal swab samples were collected from each patient, and genomic DNA was extracted from each sample. All exonic regions of all human genes (~22,000) were captured using Agilent (Santa Clara, CA, USA) SureSelect kits (version C2, December 2018) and sequenced using the NovaSeq platform (Illumina, San Diego, CA, USA). Sequence reads were aligned to the human reference sequence (original GRCh37 from NCBI, February 2009) using BWA-MEM (v.0.7.17), and variants were called following GATK germline best practice^12^.

Human phenotype ontology (HPO) terminology is used to define a symptom set of diseases^13^. All variants from a patient were counted equally for each symptom of the patient. We used OMIM as the primary source for disease identification and symptom sets in HPO terms^14^. To cover more disease–phenotype relationships, Orphanet resources are additionally used^15^.

#### Epilepsy WES data

WES data from a case-control study involving epilepsy-affected individuals were used to verify the reproducibility of our BayesianOR calculation method^16^. To obtain symptom sets from diseases, we only focused on severe developmental and epileptic encephalopathies (DEEs, MIM:308350) and genetic generalized epilepsy (GGE, MIM:600669), for which it was possible to obtain one explicit OMIM ID. Each epilepsy type had allele counts (AC) and allele numbers (AN) for the case and control groups from a total of 1,844,644 variants. The numbers of symptoms of DEE and GGE we used were 17 and 3, respectively, and HP:0002123 as the shared symptom (Supplement table. 1; The shared symptom is highligted in red.). AC and AN were counted equally for their symptoms.

#### In-house WES data and GnomAD data for the case-control study

To apply BayesianOR, the observed variants were counted equally for all shared symptoms among 2,522 in-house rare disease patients. A total of 2,804,629 variants and 1,336 symptoms were collated. Variants whose reference alleles were not found in GnomAD were excluded from Bayesian OR calculations. Finally, we obtained each AC and AN from a unique 214,434,494 pair of variants and symptoms. GnomAD v2.1.1 was used for the control group to calculate BayesianOR. Total AC and AN were taken from all variants filtered with the status ‘PASS’ or ‘AC0’ (AC = 0 and AN = 0 variants were excluded).

#### ClinVar variants

For comparisons of pathogenic variants, we classified variants into three groups based on 5 categories from ACMG/AMP guidelines^1^ “Pathogenic” or “Likely pathogenic” variants as “Pathogenic”; “Benign” or “Likely benign” as “Benign”; “Uncertain significance” as “Variant of uncertain significance”(VUS). We collected variants for each group from all ClinVar^9^ RCVs that had either OMIM or Orphanet IDs. For both “Benign” and “Pathogenic” groups, only variants with 2–4 stars were used, and for the “VUS” cohort, variants with more than 2 submitters were collated. According to the condition, 14,692 “Pathogenic” variants, 8,288 “Benign” variants, and 5,247 “VUS” variants were obtained out of a total 28,227 variants for 1,467 diseases. In these sets, variants with symptoms that were not observed from in-house data, or where the variant itself has never been observed, were excluded. In all, we used 55 variants, 1,246 variants, and 395 variants of the “Pathogenic”, “Benign”, and “VUS” categories, respectively, to examine 549 diseases. ClinVar data were downloaded from; https://ftp.ncbi.nlm.nih.gov/pub/clinvar/xml/ClinVarFullRelease_00-latest.xml.gz, which was updated in June 2020.

### Statistical analysis

All the statistical tests including Shapiro-Wilk test for the normality test prior to Mann-Whitney test(U test) and Kruskal-Wallis test(KW test) for non-parametic comparisons, and the Pearson correlation coefficient *r* for measuring the reproducibility are done by using Scipy (http://www.scipy.org/). As default, statistical significance was set at P=0.05.

## Results

To demonstrate the validity of our BayesianOR and the effectiveness of classification by this method, we examined three approaches. Firstly, we attempted to verify that BayesianOR can replicate the OR of well-studied data. To this end, two subtypes of epilepsy data were analyzed^16^. Secondly, to check the validity of independent frequency data, 3billion’s proprietary inhouse data were applied to the same variants of the above study. Finally, the distribution of BayesianOR for variants of ClinVar pathogenic/VUS/benign groups were compared to show the potential of variant accumulation by decomposed symptoms in the case of PS4 criterion.

### OR reproducibility

The results of the OR and BayesianOR are compared in Figure 1. The AC and AN tables were generated from epilepsy study data following the methods by which the in-house data were generated, and then the scores were calculated.

**Figure 1.**
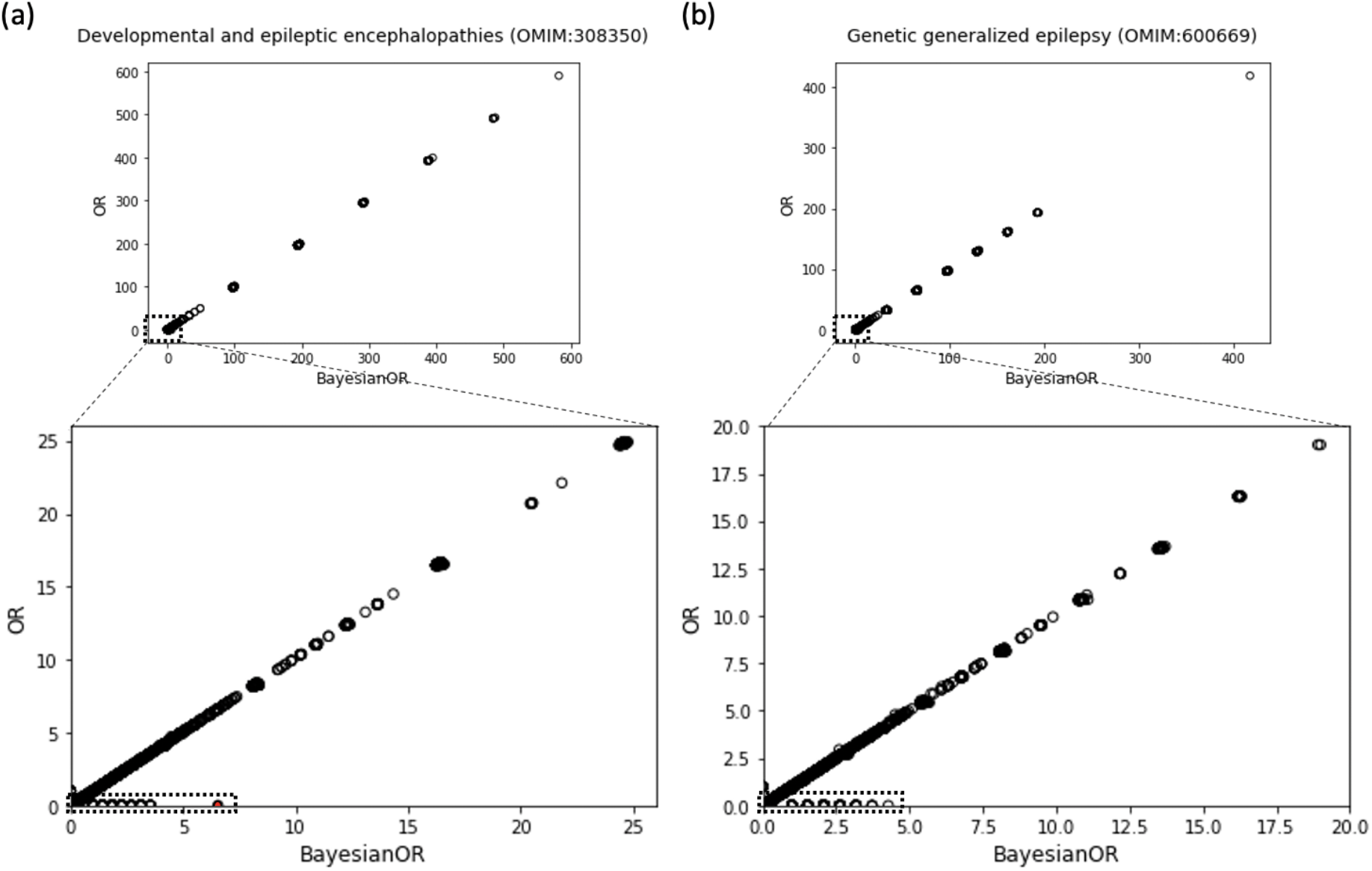
Reproducibility of OR (y-axis) and BayesianOR (x-axis). Each circle represents a variant. The high correlation between OR and BayesianOR is observed in both (a) DEE (r^2^ = 99.97) and (b) GGE(r^2^ = 99.93), respectively. All ORs, except for not observed in control data, were calculated using epilepsy study data^16^. BayesianOR could calculate the OR for some rare variants of which the original OR could not be calculated (dashed box). See text for details.

The DEE and GGE cases exhibited an almost exact correlation between the OR and BayesianOR results (DEE, r^2^ = 99.97; GGE, r^2^ = 99.93). Furthermore, due to accumulation of allele counts of the shared symptoms between DEE and GGE cases, the variant frequencies for which allele counts were not observed in cases can be calculated. For example, in the case of X-38146601-G-GT (Figure 1(a) red dot), originally, the OR could not be calculated due to zero observations in both DEE case and control populations; however, BayesianOR provided the OR for this variant (Supplement table. 2)

### In-house data-based BayesianOR result

To demonstrate the utility of our novel approach under the condition of lack of data for some diseases, the BayesianOR was calculated with the epilepsy variants using in-house data, which represented around 3,000 patients with various types of rare diseases. Due to this condition, all symptoms belonging to DEE and GGE did not always have corresponding variant allele counts. The calculation of BayesianOR was performed using only those symptoms with more than one variant.

According to the ACMG/AMP guideline of PS4 Note 1, to designate a variant to PS4, OR of the variant should be > 5, and the OR confidence interval should be > 1. Applying these guidelines, the results are described in Figure 2. Despite using in-house data to calculate BayesianOR, a distinctive pattern was observed. The distribution of BayesianOR in the high OR (> 5) group was distinguishable from that of the low OR(< 1) group (Figure 2 (a); p-value = 2.6e^-209^ and (b); p-value = 1.5e^-136^. Both p-values are from *U* tests). Distributions of variants with 95% confidence intervals show more contrast. (Figure 2 (c); p-value = 3.7e^-170^ and (d); p-value = 6.8e^-7^. Both p-values are from *U* tests).

**Figure 2.**
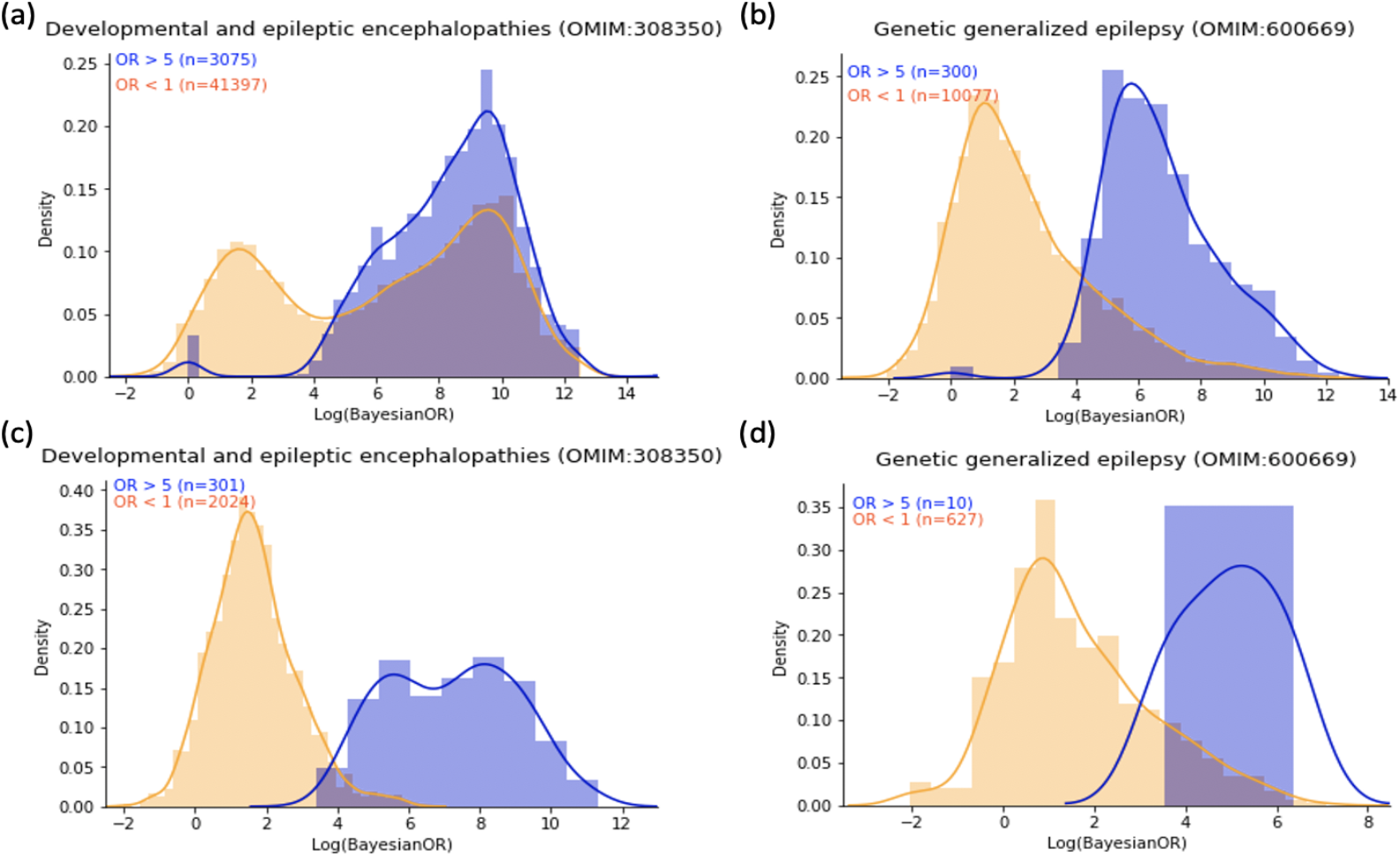
The distribution difference of BayesianOR between high OR variant (> 5) and low OR (< 1) variant groups. The calculation was performed using in-house data. Even with the difference in the accumulation of each symptom, all panels tended to show a distinctive pattern. Panels (a) and (b) represent all variants of DEE and GGE, respectively, and (c) and (d) are variants with high confidence intervals.

### ClinVar data analysis

Three categorized ClinVar variants were demonstrated, based on BayesianOR values. Indeed, all pathogenic ClinVar variants should not be categorized as a PS4 criterion, but there should be a tendency of high BayesianOR for pathogenic variants, low values for benign instances, and middle range for VUS variants. Based on our assumptions, the results depicted in Figure 3 show biased distributions. Except for a few pathogenic cases, the distributions of BayesianOR according to pathogenicity definied by ClinVar were statistically different from each other (p-value = 1.0e^-126^; KW test. Post-hoc Bonferroni corrected p-values are shown in Supplement table 3). This means that BayesianOR results calculated from in-house variant frequencies and other inferred parameters can reflect the property of pathogenicity.

**Figure 3.**
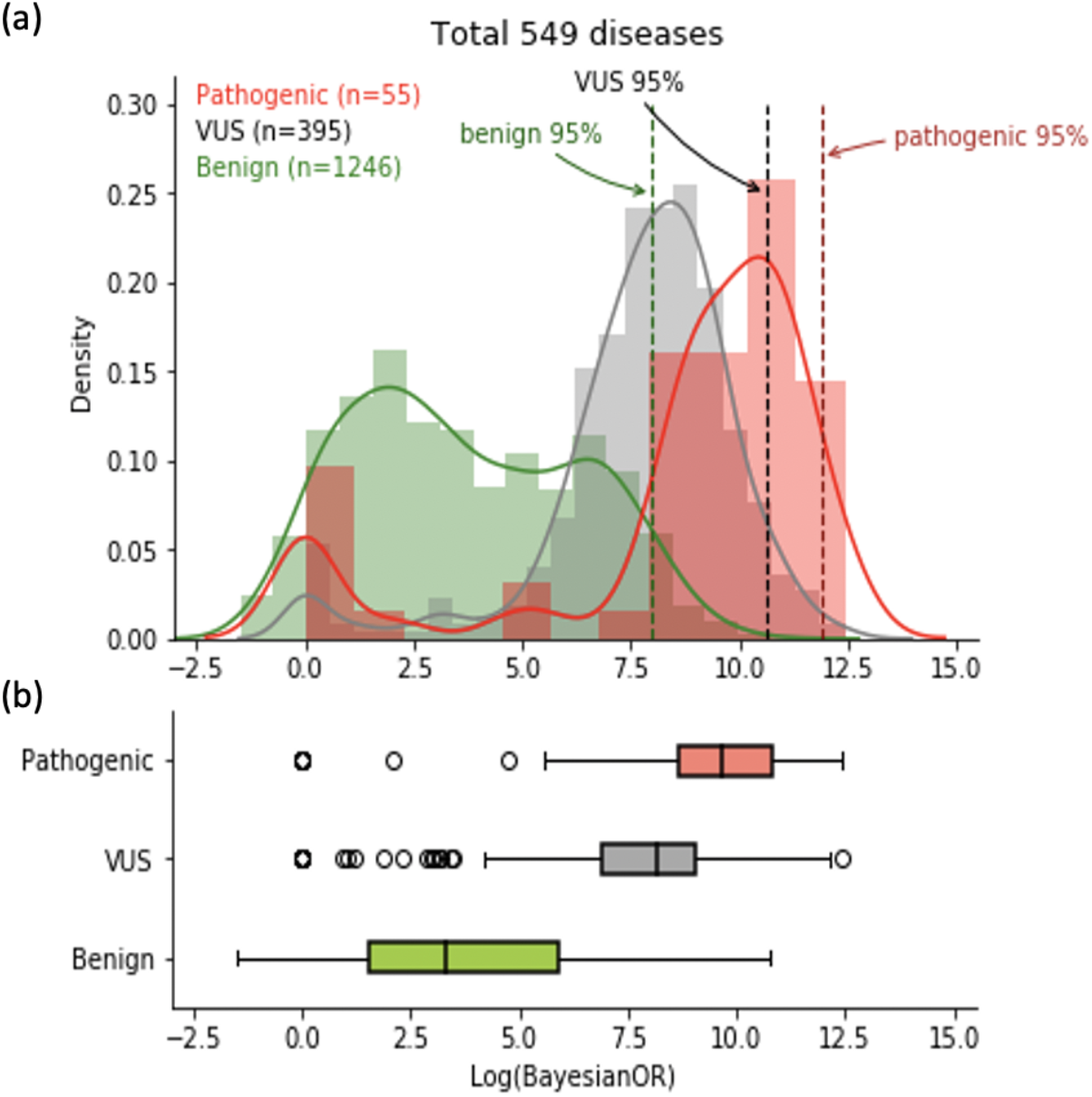
Variation of BayesianOR distribution with ClinVar pathogenicity. Pathogenic/Likely pathogenic variants are grouped as “Pathogenic” and Benign/Likely benign as “Benign”. Variants of unknown significance are indicated as VUS. (a) Histogram and density curve of BayesianOR for each pathogenicity group. (b) Boxplot of the same data as in (a).

## Discussion

The ACMG guidelines are considered a standard procedure for interpreting variant pathogenicity. Even though a strong level of evidence significantly improves the sensitivity of detection for putative disease-causing variants, it is true that applying the PS4 criterion has limitations of its own. For ultra-rare variants, expert panels such as ClinGen, have built guidelines for the criterion, but such methods take a long time to handle all kinds of variants for various rare diseases.

The novel approach of using BayesianOR of the ACMG PS4 criterion for rare variants, was proposed in this study. To evaluate the validity of our method, we checked the reproducibility of a previously studied epilepsy data set. Based on the results obtained, BayesianOR exhibited reproducible results (DEE, r^2^ = 99.97; GGE, r^2^ = 99.93). To verify the applicability of our method, we applied our in-house variant data to the epilepsy study and also obtained statistically meaningful results for high and low OR groups (DEE, p-value = 2.6e^-209^; GGE, p-value = 1.5e^-136^). For ClinVar variants, BayesianOR shows discriminate distributions of scores with in-house variant data (p-value = 1.0e-126, Kruskal-Wallis test), based on which we conclude that the Bayesian approach using decomposed symptoms can be applicable for the PS4 criterion of rare variants. Symptoms are shared among many different rare diseases, and diseases can be decomposed to many symptoms. Based on this idea, the number of allele counts of the variants can be accumulated.

The scope of this work is portrayed in Figure 4. Even though BayesianOR is calculated with a limited number of patients, the results of this research showed a certain degree of meaning for calculating ORs for the impossible cases when using the original method. If patient data with accompanying symptom information are more collated, we believe the performance of BayesianOR can reachthe extent of the original OR.

**Figure 4.**
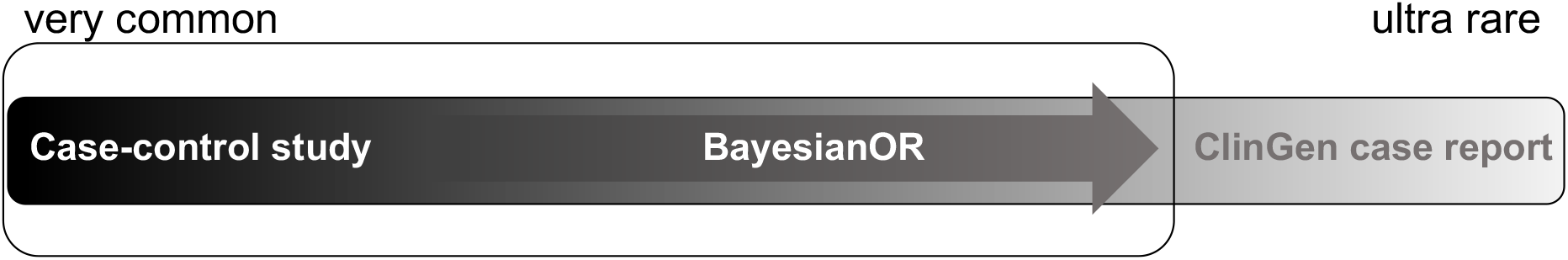
BayesianOR scope. With allele count accumulation by decomposing diseases into symptoms, the OR calculation can be extended to the rarer variants.

In applying this method to PS4 criterion, the current approach has certain limitations. Firstly, OR is calculated from a case-control study, whereas BayesianOR scores use the GnomAD database as a control group. Many previous papers have reported that GnomAD has minimal patient data, especially for late-onset diseases. Secondly, it is not easy to define the BayesianOR threshold. All calculations are performed using in-house variant data, and this makes the range of BayesianOR scores vary depending on the status of each institution’s database accumulation. Moreover, confidence intervals cannot be easily obtained for the same reason. Finally, the additive model of symptoms for each disease has the possibility to overstate the BayesianOR, rather than the real OR. A more sophisticated disease model of symptoms may be devised to reduce such inflation.

Even though PS4 is an important criterion for designating the pathogenicity of novel variants, the application of this rule is not easy owing to variant rarity. Due to this limitation, the ACMG 2015 guidelines present a second caveat regarding the number of probands, and the ClinGen expert panels have reviewed certain genes and variants as well^3–8^. Nevertheless, these efforts seems far from covering all rare diseases. In this study, a novel approach based on Bayesian framework was presented to expand the OR for many rare variants using symptom-based calculations. In the future, as more patient data accumulate, the usability of this approach can be increased widely for rare variants, whose OR cannot be otherwise calculated at present.

## Acknowledgment

Institute for Information and Communications Technology Promotion (IITP) grant funded by the Korean government (MSIT) (2018-0-00861, Intelligent SW Technology Development for Medical Data Analysis).

