## Supplemental data for "A novel PS4 criterion approach based on symptoms of rare diseases and in-house frequency data in a Bayesian framework"

**Supplement figure 1.**

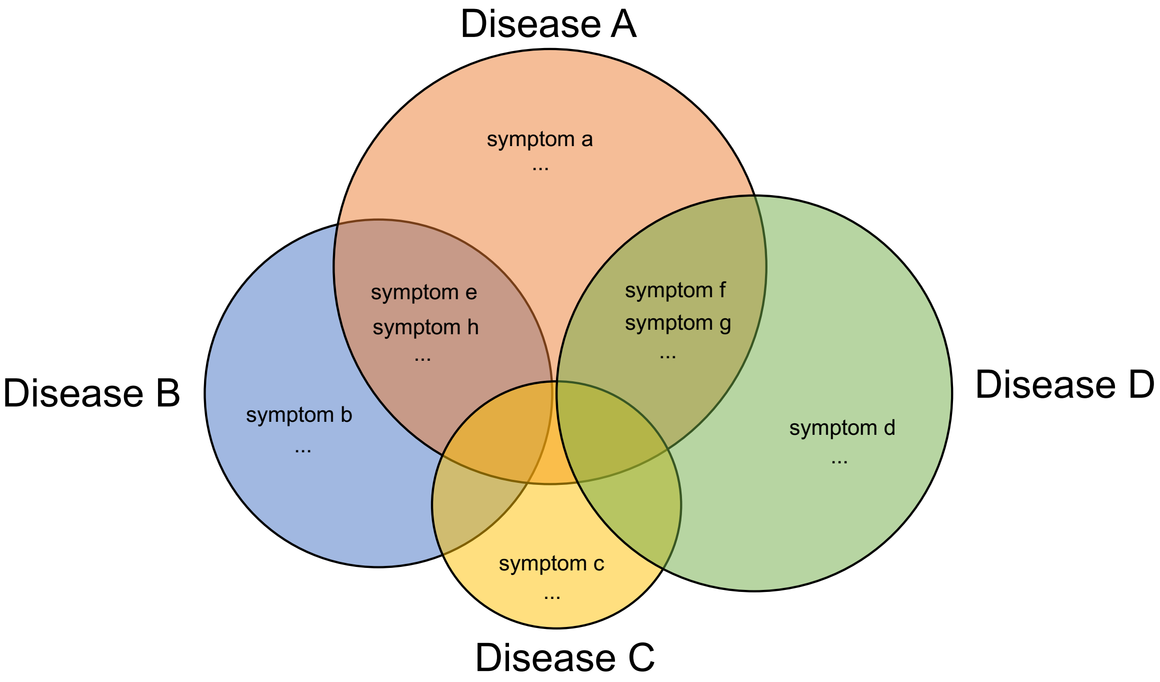


**Supplement figure 1. A diagram showing an example of sharing symptoms between diseases.**

### Supplement Table 1.

| Disease | Symptoms |
|---|---|
| DEE | HP:0011097, HP:0002098, HP:0001419, HP:0001332, HP:0200136, HP:0001266, HP:0002015, HP:0008936, HP:0001257, HP:0001249, HP:0040195, HP:0002119, HP:0002123, HP:0001347, HP:0002521, HP:0002094, HP:0001276, HP:0000006 |
| GGE | HP:0002121, HP:0002123, HP:0002069 |

### Supplement Table 2.

| Phenotype | AC | AN |
|---|---|---|
| HP:0001249 | 0 | 2042 |
| HP:0001257 | 0 | 2042 |
| HP:0001266 | 0 | 2042 |
| HP:0001276 | 0 | 2042 |
| HP:0001332 | 0 | 2042 |
| HP:0001347 | 0 | 2042 |
| HP:0001419 | 0 | 2042 |
| HP:0002015 | 0 | 2042 |
| HP:0002094 | 0 | 2042 |
| HP:0002098 | 0 | 2042 |
| HP:0002119 | 0 | 2042 |
| HP:0002123 | 13 | 8256 |
| HP:0002521 | 0 | 2042 |
| HP:0008936 | 0 | 2042 |
| HP:0011097 | 0 | 2042 |
| HP:0040195 | 0 | 2042 |
| HP:0200136 | 0 | 2042 |

### Supplement Table 3

| U test | Pathogenic vs. VUS | Pathogenic vs. Benign | Benign vs. VUS |
|---|---|---|---|
| adjusted p-value | 3.39E-07 | 4.41E-20 | 2.33E-117 |
| statistics | 6185 | 9120 | 56947 |
